# Comparison of early effects of PCV7, PCV10 and PCV13 on *S. pneumoniae* (SP) nasopharyngeal carriage in a population based study; the Palestinian-Israeli Collaborative Research (PICR)

**DOI:** 10.1101/409359

**Authors:** Rania Abu Seir, Kifaya Azmi, Ayob Hamdan, Hanan Namouz, Fuad Jaar, Hanaa Jaber, Carmit Rubin, Dafna Doron, Galia Rahav, Ziad Abdeen, Gili Regev-Yochay

## Abstract

**Background:** Pneumococcal conjugate vaccines (PCVs), PCV10 and PCV13, are currently used in different countries. We have previously reported the effectiveness of PCV7, following its introduction in Israel and before PCVs were introduced in Palestine. Here, we extended the study and compared the initial impact of PCV10 to that of PCV7/13.

**Methods:** Four cross-sectional surveys of *S. pneumoniae* carriage among children <5y through 2009-2014 were preformed among two proximate populations, living under two distinct health authorities, with different vaccination policies. In East-Jerusalem (EJ), PCV7 was implemented in 2009 and replaced by PCV13 in late 2010, while in Palestine (PA), PCV10 was implemented in 2011.

**Results:** A total of 1267 and 2414 children from EJ and PA were screened. Implementation of both PCV7 (in EJ) and PCV10 (in PA) did not affect overall *S. pneumoniae* carriage (∼30%), but resulted in a significant decrease in carriage of VT7 strains. In the pre-vaccine era, VT7/VT13 strains consisted 47.0%/62.0% and 41.2%/54.8% of pneumococci in EJ and PA, respectively. A 48.6% and 53.9% decrease was observed within 3 years of PCV7 implementation in EJ (p= 0.001) and PCV10 in PA (p<0.0001), respectively. These vaccination policies also resulted in ∼50% reduction in VT13-added serotypes especially 6A (from 11.0% to 0.0% (EJ) and 9.5% to 4.9% (PA)). Three years after PCV13 implementation in EJ, an additional 67% decrease in VT13 strains was observed, yet an increase in serotype 3 was observed (0.0% to 3.4%, p=0.056). The prevalence of non-VT13 strains increased during the study period from 38% and 45.3% to 89.8% and 70.7%, in EJ and in PA respectively.

**Conclusions:** Within the first three years following PCV implementation, we observed similar reductions in carriage of VT10 and VT13 strains with either vaccination policies, with no effect on overall carriage. Further follow-up is needed to compare the long-term effects.

## Introduction

Pneumococcal diseases cause high rates of morbidity and mortality worldwide causing bacterial meningitis, community-acquired pneumonia, acute otitis media, and sinusitis [1]. Nasopharyngeal (NP) colonization with *S. pneumoniae* is common in young children and serves as the source of transmission between individuals in the community and as the first step towards infection [2]. Moreover, understanding the dynamics of pneumococcal colonization following PCV implementation in children will lead to better understanding and possible prediction of the herd effects associated with PCV implementation [3]. The introduction of pneumococcal conjugate vaccine (PCV) to the routine childhood vaccination had a dramatic impact on the incidence of invasive pneumococcal disease (IPD) due to decrease in vaccine-type (VT) colonization and infections [4]. Currently two PCVs are in use in different countries; PCV10 is being used in several South American countries, Finland, the Netherlands and in Palestine, covering serotypes 1, 4, 5, 6B, 7F, 9V, 14, 18C, 19F and 23F. PCV13 is being used in the USA, most European countries, several African countries, Australia and Israel [5]. PCV13 covers all PCV10 serotypes and additionally, serotypes 3, 6A and 19A. Despite the greater number of serotypes covered by PCV13, it is still controversial whether its impact on carriage and disease are greater [6].

The Palestinian-Israeli Collaborative Research (PICR) group was established in 2009, by independent Palestinian and Israeli researchers, and offers a golden opportunity to study vaccination effects and impacts in the region, where two closely related Palestinian populations are governed by two distinct health authorities with different vaccination policies [7]. In this study, we compare the effects of sequential PCV7/PCV13 to PCV10 implementation on Palestinians living in East Jerusalem under Israeli health policy, vaccinated with PCV7/13 and Palestinians living in the West Bank, under the Palestinian Authority health policy, vaccinated with PCV10.

## Materials and methods

### Ethical approvals

The study was approved by the Institutional Review Boards (IRB) of Sheba Medical Center, Maccabi Healthcare Services (MHS) and Al-Quds University. The parent of each participating child provided a written informed consent before recruitment.

### PICR setting and study population

A population based study consisting of repeated cross-sectional surveillances was conducted in two geographically proximate Palestinian populations which are under two different health authorities and consequently under different vaccination policies. This setting was previously described in detail [7]. In brief, Palestinian children (<5 years) and their accompanying parent who visited their primary care pediatrician, either from East Jerusalem (EJ), which is under Israeli Health Law, or from Palestinian cities in the West Bank governed by the Palestinian Authority (PA) were enrolled if the parent agreed and signed an informed consent.

### Vaccination policies in the two populations

In Israel, PCV7 was approved for use and introduced to the private market in 2007 and to the pediatric national immunization program (NIP) in July 2009 in a 2+1 schedule with a 2 dose catch-up. In November 2010, PCV13 gradually replaced PCV7 without catch-up. In Palestine, until 2011, access to PCV was very limited. In 2011, PCV10 was introduced through the Palestinian Ministry of Health to the pediatric NIP in a 2+1 schedule through primary care physicians. The vaccination was provided free of charge to all children under 2y by the Palestinian Ministry of Health.

### Study design and pneumococcal carriage screening

Four cross-sectional surveillance studies were conducted during May-July of 2009-2011 and 2014, results of three which have been previously reported [7]. The first surveillance, in 2009, served as a baseline, pre-vaccination surveillance for both populations, before PCV was introduced in either PA or EJ. The following two surveillances were conducted in the consecutive years (2010-2011) and allowed to evaluate the impact and effects of PCV7 [7]. This update of the study includes the forth surveillance, conducted in 2014, three and a half years after PCV13 replaced PCV7 in EJ and three years after the introduction of PCV10 in PA. This surveillance provided the opportunity to assess the initial effects of PCV10 compared to PCV13. Children and their parents were screened for nasopharyngeal pneumococcal carriage as previously described [7]. In brief, medical history data, including vaccination history, were collected from the parents by a study coordinator and from the medical files by the physicians. Swabs were transferred to the laboratory within 24 hours, where pneumococci were identified and serotype was determined in two steps; initially latex agglutination test (StatenSerum Institute, Copenhagen, Denmark) was used to determine the serogroup, followed by PCR to determine the serotype.

### Statistical analysis

Prevalence rates and proportions were calculated and compared using Chi-square or Fisher’s exact test, as applicable. The Cochran-Armitage trend test was used to determine trends along the study years. Additionally, comparison of VT13 prevalence between the two regions before and after the implementation of the three different vaccines was applied using the Cochran–Mantel– Haenszel test. All statistical analyses were performed using SAS 9.4 for Unix.

## Results

### Study population description

A total of 1267 and 2414 children from EJ and PA were screened in the four surveillance studies [7]. Each year, approximately 300 children were screened in EJ and 600 children were screened in PA. A detailed description of the study population of the first three surveys (2009-2011) was previously reported [7, 8]. Here, we present an additional surveillance that took place in 2014 in which 287 children from EJ and 643 children from PA were screened. Population characteristics in this surveillance are similar to those in the 3 initial surveillances, with a slightly higher proportion of males (54% and 59% in EJ and PA), approximately, 45% of the children between 6 and 24 months and >15% of the children living in large households with 7+ household members, in both regions (S1 **Table**).

### PCV coverage

During the first year of the study, in the pre-vaccine period, when PCV7 was approved and available in the private market, but not yet implemented in the NIP, we found that only 2.7% and 2.3% of the children in EJ and PA respectively received at least one dose of the vaccine, and only 1.2% in EJ and 0.7% in PA received at least two doses. Within a year of implementing PCV7 in EJ, the proportion of children <2 years old who received at least one vaccine dose increased to 75%. In PA, PCV10 was not introduced until late 2011, the proportion of children <2 years who received at least one dose of any PCV (PCV7 or PCV10) in PA before PCV10 implementation did not exceed 13%. This proportion increased to 85.5% three years after implementing PCV10 in the vaccination program compared to 92% of children in EJ (p=0.03) **(Table 1).**

**Table 1:**
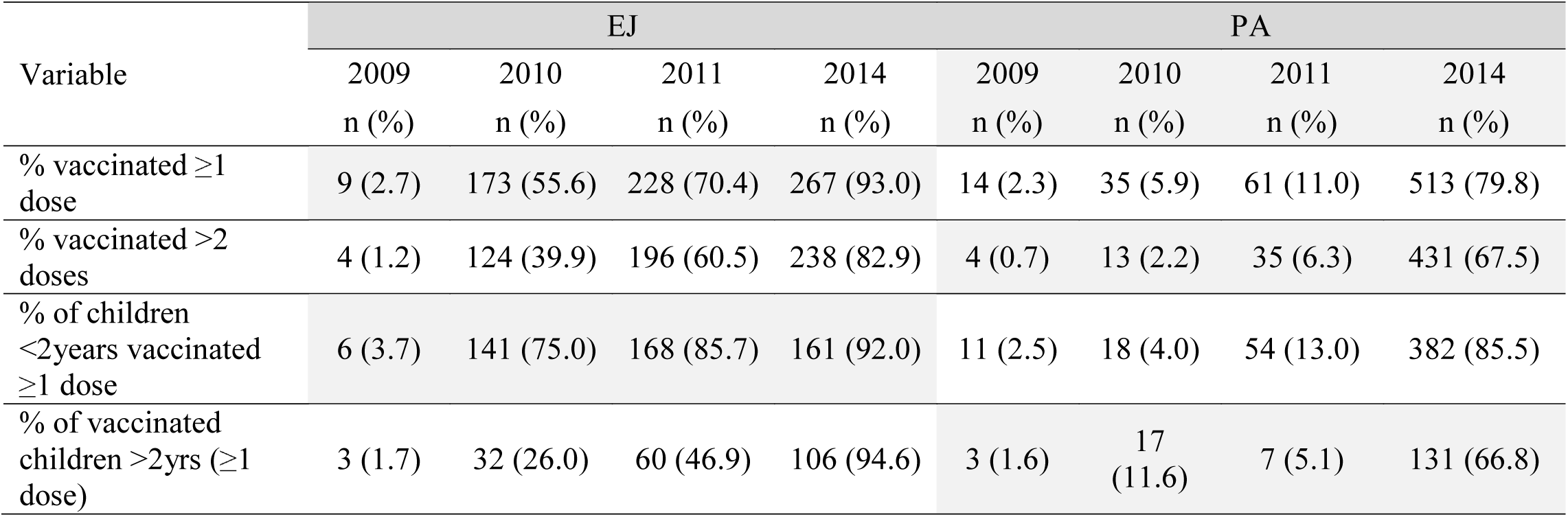
Vaccine coverage in study population by year and region.

### *S. pneumoniae* carriage and serotypes’ distribution

Of all children screened, *S. pneumoniae* carriage was detected in 29.0% (100/345) of the children in EJ and 36.0% (223/620) of children in PA in 2009 before PCV was introduced. This carriage did not change significantly in EJ during the study period and ranged between 26.9% and 30.7%. In PA, *S. pneumoniae* carriage decreased from 36.0% (223/620) to 28.8% (160/556) (p_trend_=0.0092) during the first three study years (before PCV10 introduction), but did not decrease further, three years after PCV10 was introduced **(Fig 1).** The overall *S. pneumoniae* carriage among children during the whole study period was 28.9% in EJ and 31.8% in PA.

**Fig 1.**
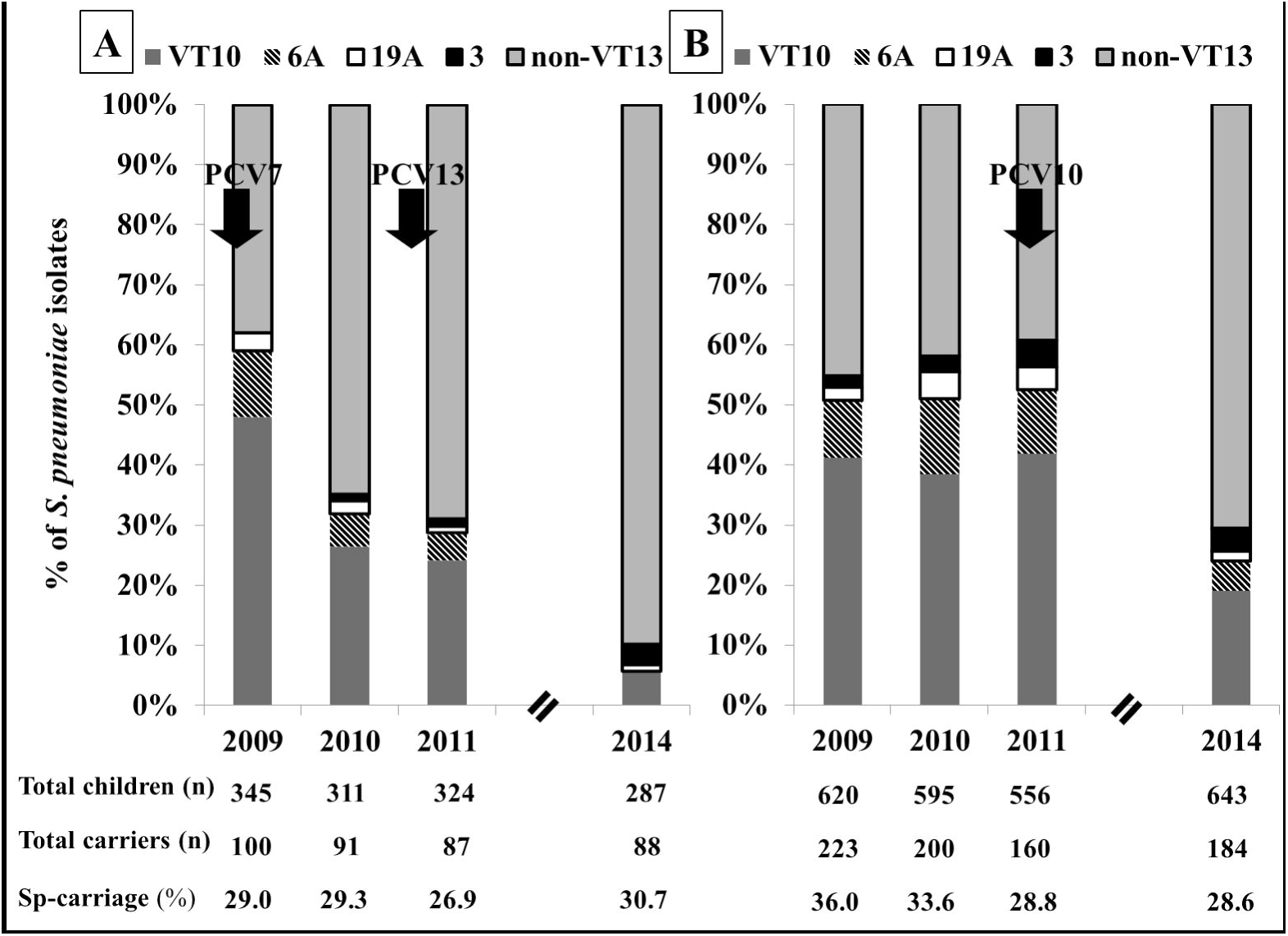
Proportion of VT strains’ carriage among all *S. pneumoniae* isolates in the two populations. (A) In EJ. (B) In PA. Dark grey represents the proportion of VT10 strains among all isolates. Stripes represent the proportion of carriage of 6A serotype. White represents the proportion of carriage of 19A serotype. Black represents the proportion of carriage of serotype 3, and Light grey represents the proportion of carriage of non-VT13 serotypes.

In the pre-vaccine study year, the proportion of VT strains among all pneumococcal isolates was similar in the two populations, with VT7 strains constituting 47.0% of all isolates in EJ and 41.2% in PA (p=0.33). VT10-added serotypes (1, 5, and 7F) were rarely carried (3/1130) during all years in both populations. Prior to vaccine introduction in 2009, VT13-added serotypes (3, 6A and 19A) constituted 14.0% (n=14) in EJ and 13.6% (n=30) in PA and non-VT13 serotypes constituted 38.0% (n=38) of all *S. pneumoniae* in that year in EJ and 45.3% (n=100) in PA. Since the VT10-added serotypes were very rarely carried, the impact on VT7 carriage was identical to that on VT10; therefore, we report only the impact on VT10 and VT13. During the first two years following PCV7 implementation in EJ, but not yet in PA, a significant decrease in VT10 strains was observed in EJ (p=0.0007), while no change was observed in PA (p=0.8914), as described previously [7].

A further significant decrease in VT10 serotypes was observed three years after the introduction of PCV13 in EJ, with prevalence of VT10 strains decreasing to 5.7% (n=5) in 2014 (p=0.0006). In PA, following introduction of PCV10 in 2011, the prevalence of VT10 serotypes significantly decreased from 41.9% (n=67) in 2011 to 19.0% (n=35) by 2014 (p<0.0001) **(Fig 1).**

Concurrently, carriage of non-VT13 serotypes, increased over the years after introducing PCV7/13 in EJ and they gradually replaced VT13 serotypes. In PA, carriage of non-VT13 serotypes did not change before PCV10 introduction (p=0.2527), but 3 years later the prevalence almost doubled, replacing VT13 serotypes (p<0.0001) **(Fig 1).**

When we assessed the rate of decrease in VT13 strains following the introduction of each of the three vaccine introductions, a similar 50% reduction was observed within 3 years of implementation of either PCV7 or PCV10 and an additional reduction of 67% in VT13 strains was observed after PCV13 introduction **(Table 2).** Overall, under the sequential PCV7/13 policy, VT13 strains decreased by 83.5% during the whole study period (from 62.0% to 10.2%) in EJ, but during the first two years following PCV7 implementation VT7 strains decreased by 48.7% (from 47.0% to 24.1%), and following PCV13 implementation, within 3.5 years VT13 strains decreased by another 67.1%. In PA, where PCV10 was implemented three years later, VT10 strains decreased by 54.7% (from 41.9% in 2011 to 19.0% in 2014) but interestingly the VT13- added serotypes also decreased by 45% (p=0.0258), this decrease was attributed to the decrease in serotype 6A **(Table 2).**

**Table 2:**
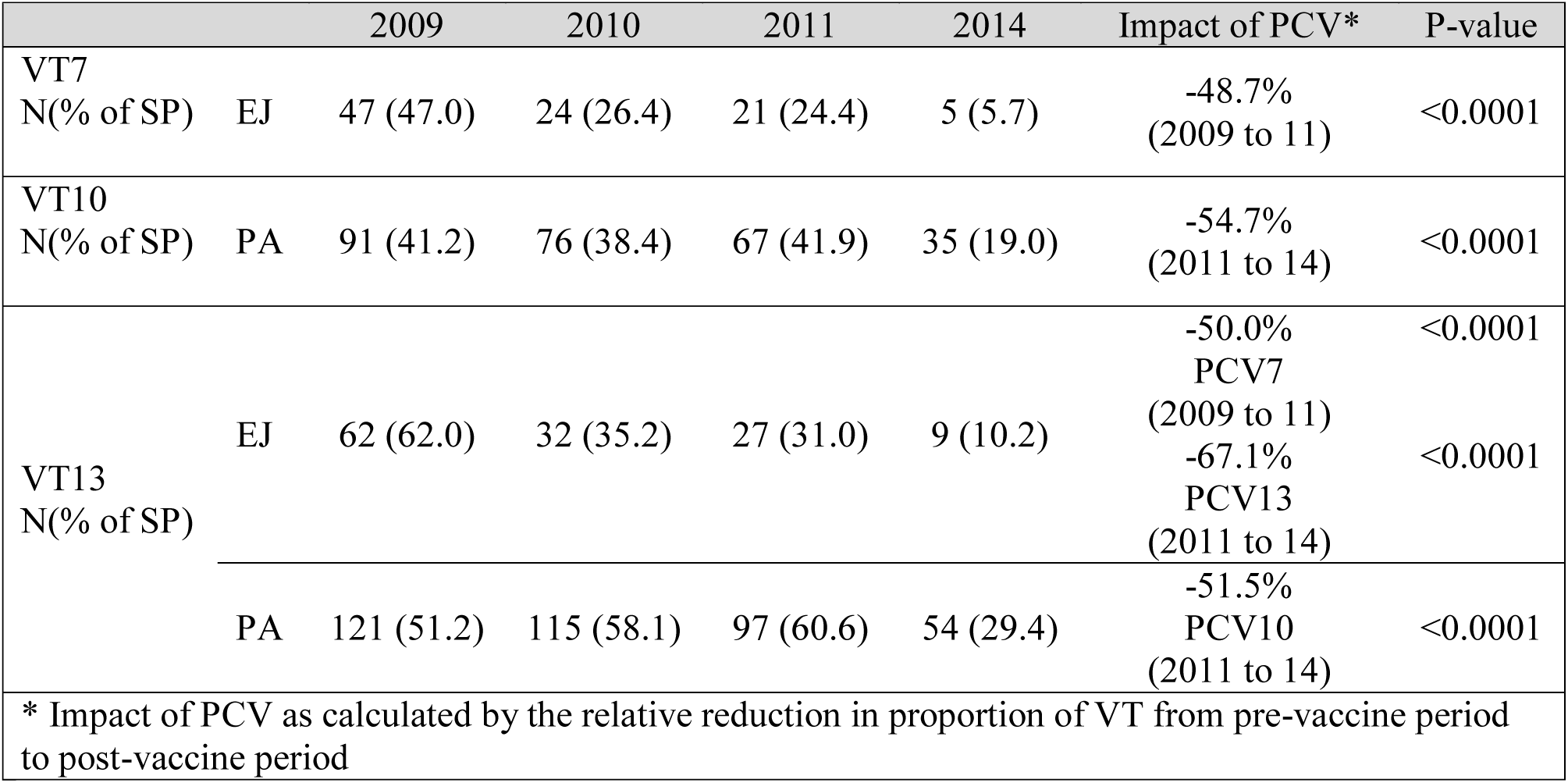
Proportion of VT13 decrease post PCV7/10/13 introduction.

### Serotype distribution and replacement

While VT10 serotypes decreased significantly following implementation of any of the vaccines, these were replaced by non-VT13 serotypes. In EJ, VT10 serotypes were nearly eliminated by 2014 with only a few isolates of serotypes 14, 23F, and 9V remaining. Yet, the group of VT13- added serotypes (3, 6A and 19A) did not decrease significantly in EJ after the introduction of PCV13 (p=0.536). The dynamics of each of these serotypes was different. Serotype 6A decreased significantly, already following PCV7 introduction, and was completely eliminated by 2014. Similarly, serotype 19A decreased somewhat (not statistically significant) following PCV7, but no further decline was observed following PCV13 introduction. However, at the same time serotype 3 increased from 0 cases in 2009 to 3/88 (3.4%) in 2014, although this increase was only nearly statistically significant (p=0.056) **(Table 3)**. In PA, before PCV10 implementation, a relatively stable prevalence of VT10 and VT13-added serotypes was observed. Within three years following PCV10 implementation, a significant decrease in VT10, which was attributed to large decrease in serotypes 6B, 23F and 19F, was observed (from 30.0% from the carriage in 2009 to 9.2% in 2014). Interestingly, a decrease was also observed in the prevalence of two of the VT13-added serotypes: 6A (from 10.6% (n=17) to 4.9% (n=9), p=0.045) and 19A (from 3.8% (n=6) to 1.6% (n=3), though not statistically significant). The prevalence of serotype 3 did not change in this period (∼4%).

**Table 3:**
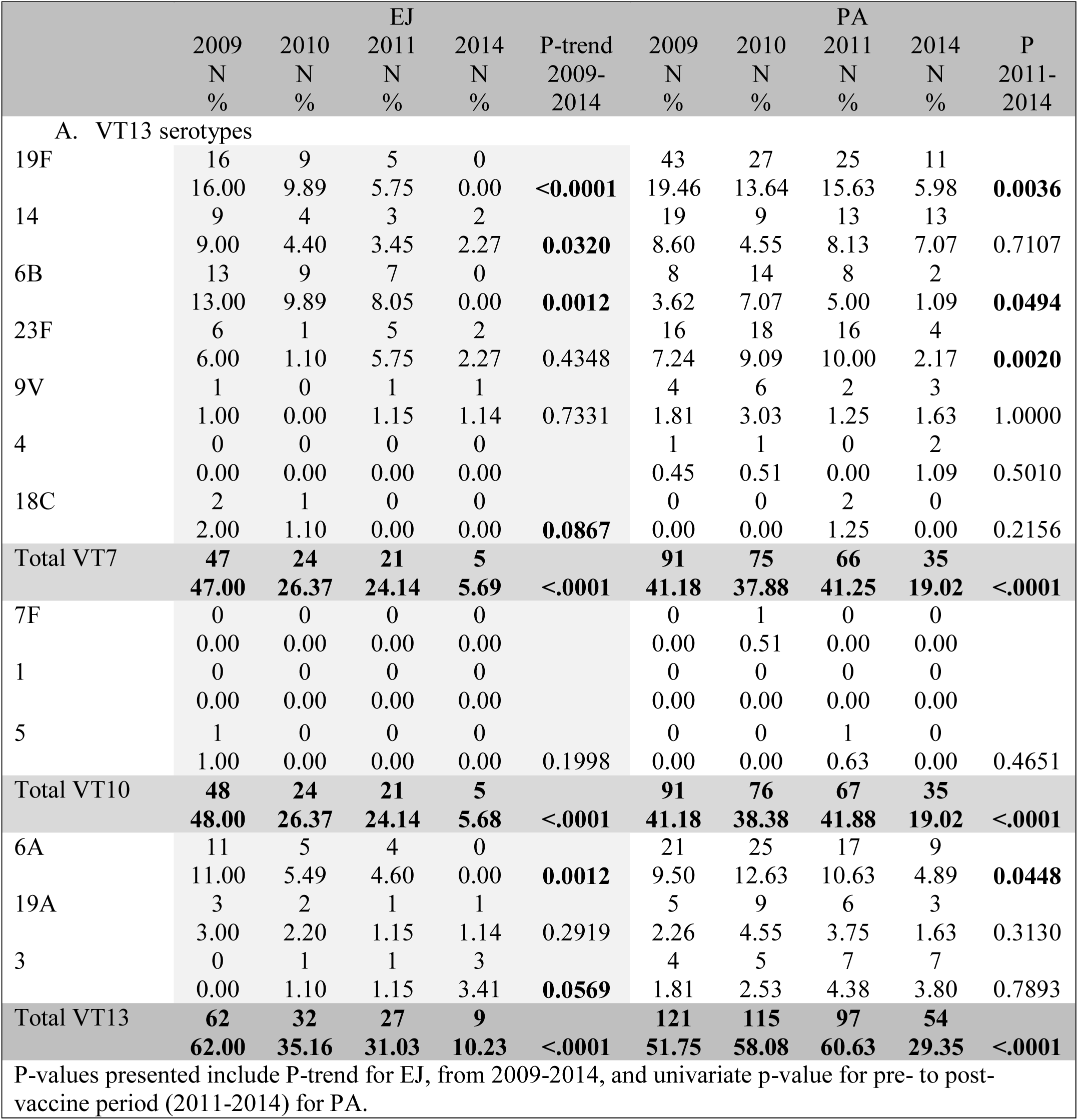
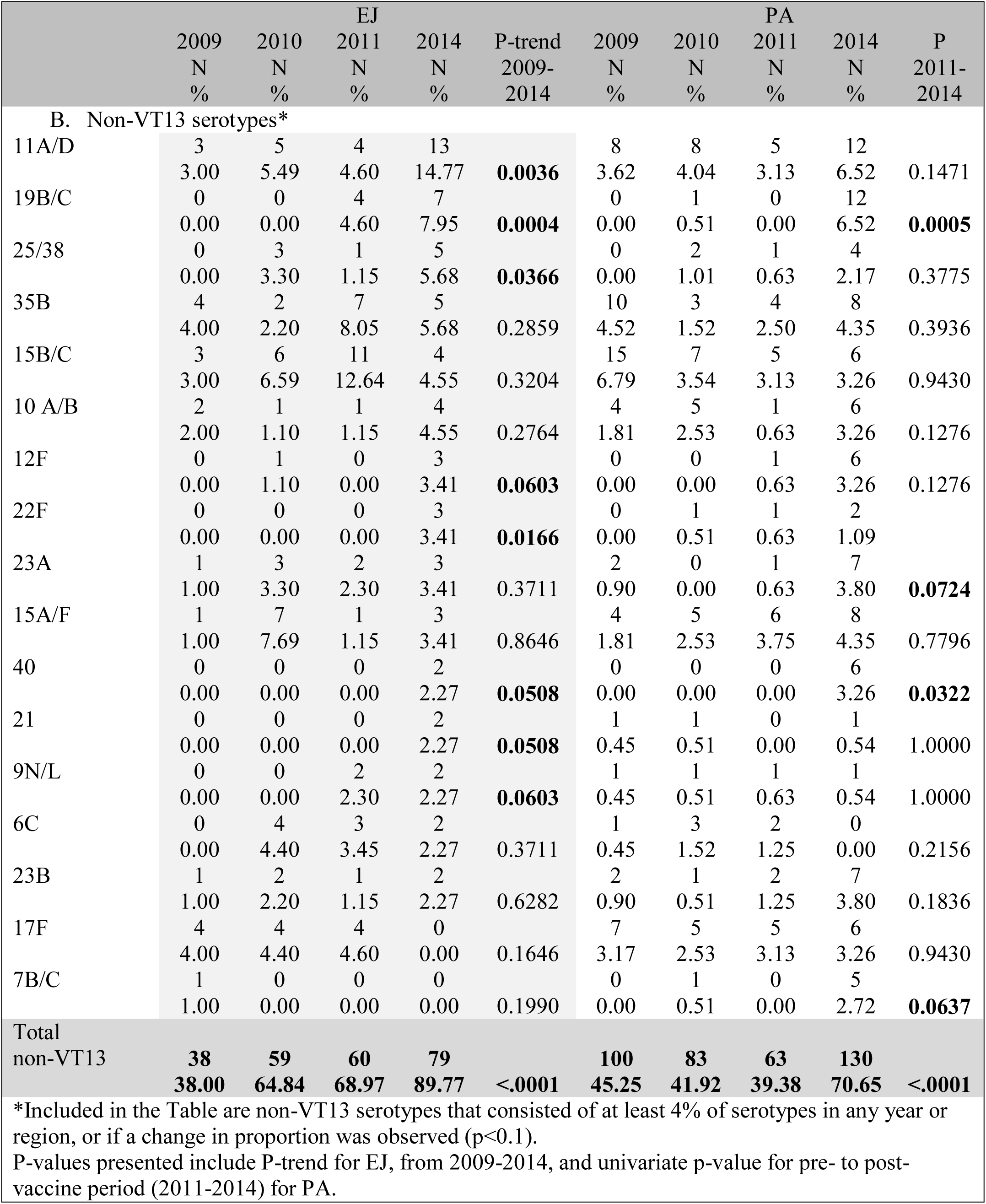
Prevalence of *S. pneumoniae* serotypes in the two regions by year.

P-values presented include P-trend for EJ, from 2009-2014, and univariate p-value for pre- to post- vaccine period (2011-2014) for PA.

*Included in the Table are non-VT13 serotypes that consisted of at least 4% of serotypes in any year or region, or if a change in proportion was observed (p<0.1). P-values presented include P-trend for EJ, from 2009-2014, and univariate p-value for pre- to post- vaccine period (2011-2014) for PA.

Non-VT13 serotypes gradually replaced VT13 serotypes; therefore, overall carriage of *S. pneumonia* did not change significantly in both regions. The proportion of non-VT13 strains increased from 38.0% of all isolates in pre-PCV surveillance in 2009, to 89.8% three years after the introduction of PCV13 in EJ (p<0.001). As for PA, the proportion of non-VT13 serotypes increased from 39.4% in the pre-PCV period (2011) to 70.7% in 2014 (p<0.001). **Table 3** presents the most common non-VT13 serotypes in both regions and their proportion among all isolates during the four surveillance periods, as well as p-values of the change pre- to post-vaccine periods. By 2014, serotypes 19B/C, 11A/D, 22F, 25/38, 40, 21, 9N/L and 12F emerged in EJ, while in PA, the non-VT13 serotypes that emerged 3 years following PCV10 implementation were 19B/C, 40, 23A and 7B/C.

### Parental carriage

*S. pneumoniae* carriage among the parents was relatively rare, with 3.8% (n=139/3681) of parents detected as nasopharyngeal carriers in both regions. Overall, 18.3% (n=23) of parent strains belonged to VT10 serotypes in both regions. Serotypes 14, 19F and 23F constituted the majority of VT10 serotypes (73.9%, n=17). Parental strains that belonged to non-VT13 serotypes constituted 67.5% (n=27) and 72.1% (n=62) in EJ and PA, respectively **(Table 4).** The small sample size of parental strains did not allow us to assess PCV effect on parental carriage or strain distribution. Sixty percent of the parents who were carriers, had a child who was also a pneumococcal carrier, yet, only 42.7% of those parent-child co-carrier pairs had an identical serotype on screening. In both regions, once PCV was implemented, none of the VT13 carrier parents had a child who carried a VT13 strain.

**Table 4:**
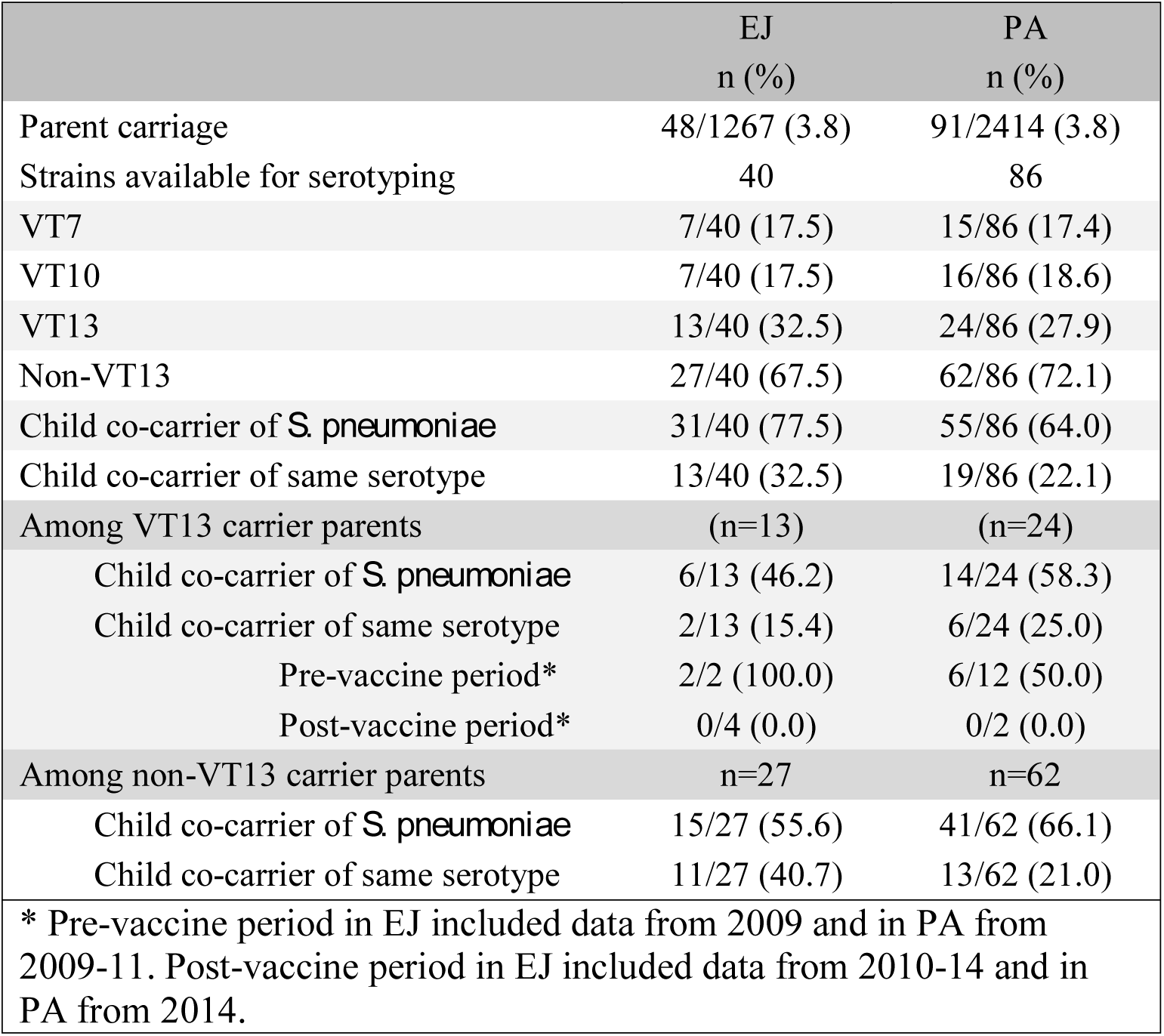
*S. pneumoniae* carriage among parents.

|  | EJ<br>n (%) | PA<br>n (%) |
| --- | --- | --- |
| Parent carriage | 48/1267 (3.8) | 91/2414 (3.8) |
| Strains available for serotyping | 40 | 86 |
| VT7 | 7/40 (17.5) | 15/86 (17.4) |
| VT10 | 7/40 (17.5) | 16/86 (18.6) |
| VT13 | 13/40 (32.5) | 24/86 (27.9) |
| Non-VT13 | 27/40 (67.5) | 62/86 (72.1) |
| Child co-carrier of *S. pneumoniae* | 31/40 (77.5) | 55/86 (64.0) |
| Child co-carrier of same serotype | 13/40 (32.5) | 19/86 (22.1) |
| Among VT13 carrier parents | (n=13) | (n=24) |
| Child co-carrier of *S. pneumoniae* | 6/13 (46.2) | 14/24 (58.3) |
| Child co-carrier of same serotype | 2/13 (15.4) | 6/24 (25.0) |
| Pre-vaccine period* | 2/2 (100.0) | 6/12 (50.0) |
| Post-vaccine period* | 0/4 (0.0) | 0/2 (0.0) |
| Among non-VT13 carrier parents | n=27 | n=62 |
| Child co-carrier of *S. pneumoniae* | 15/27 (55.6) | 41/62 (66.1) |
| Child co-carrier of same serotype | 11/27 (40.7) | 13/62 (21.0) |
| * Pre-vaccine period in EJ included data from 2009 and in PA from 2009-11. Post-vaccine period in EJ included data from 2010-14 and in PA from 2014. |  |  |

## Discussion

We have previously reported the effects of PCV7 by comparing two closely related populations in which PCV7 was implemented in one but not yet in the other [7]. In this study, we conducted an additional surveillance following the introduction of PCV10 to the previously unvaccinated population. This allowed us to assess the impact of different PCVs on pneumococcal carriage among children and their parents and the effects of the different vaccination programs. To overcome seasonal variability, all surveillances took place during spring-summer. This could explain the relatively low carriage rates on one hand, but assured that the changes observed were not confounded by seasonality.

The main vaccine impact of PCV implementation was the significant decrease in VT serotypes. More so, we show that a reduction of ∼50% in VT13 serotypes was observed within three years of implementation of PCV, whether PCV7 or PCV10 were used. This reduction was similar, despite slightly different vaccine coverage rates (with 85% vaccine coverage of <2y in PA vs. 92% coverage in EJ). Comparing the impact of PCV13 is only partially accurate, since PCV13 was introduced after a significant impact of PCV7 was observed. Yet, an additional 67% decrease in VT13 was observed 3 years after PCV13 replaced PCV7.

There is controversy over the advantage of PCV13 compared to PCV10. While some studies reported higher effectiveness and cost-effectiveness of PCV13 compared to PCV10 [9-11] due to its greater coverage, one study reported 97% effectiveness against VT-IPD for PCV10, with only 86% effectiveness for PCV13 [12] and other studies [13-15] showed no differences between them. The disparities in results of different studies, particularly regarding the effect on 19A, which was shown to emerge following PCV10 in some studies but decrease, in other studies, could be attributed to differences in the study design, different outcomes assessed (carriage, IPD, etc.), or different vaccine coverage. Alternatively, the differences could be due to differences in background circulating clones in the different geographic regions.

Particularly interesting is the comparison of the impact of the different PCVs on serotypes covered by one vaccine, but not the other (serotypes 3, 6A and 19A). Serotype 6A has been repeatedly reported to decrease following PCV7 or PCV10 implementation [14, 16-20], probably due to cross-protection by 6B in PCV7/PCV10. Serotype 19A, which is not included in either PCV7or PCV10, could similarly be cross-protected by 19F which is included in both these vaccines. However, only modest protection has been suggested [21]. Moreover, PCV7 introduction in many countries, led to emergence of serotype 19A both in carriage and in IPD [22-24], yet, we did not observe emergence of 19A in EJ after PCV7 and before PCV13 was implemented. Similarly a nationwide IPD surveillance study in Israel did not report 19A increase following PCV7 introduction among children or adults [25, 26]. A plausible explanation could be the rapid transition from PCV7 to PCV13 within less than 2 years in Israel, or a different clonal background distribution.

In contrast to PCV7, the impact of PCV10 on serotype 19A is much more controversial. Several studies reported PCV10 effectiveness against serotype 19A [12, 13, 27], while other studies reported emergence of serotype 19A following PCV10 introduction [14, 28-31]. Here, we report that serotype 19A did not emerge, or rather tended to decrease following PCV10 implementation in PA. However, it is important to note that these are only short-term (three years) observations, and longer follow-up is required to determine the long-term impact of PCV10 on serotype 19A. The last of the three additional serotypes not included in PCV10 is serotype 3. Serotype 3 is unique in many aspects, heavily encapsulated with a mucoid phenotype, highly resistant to phagocytosis. While some have suggested that despite this it is not invasive [32], others have reported it to be highly invasive, with high case fatality ratios, particularly in adults [33, 34].

Many studies from different geographical regions have reported ineffectiveness of PCV13 against serotype 3 [14, 35-38], although a few have reported decrease in serotype 3 following PCV13 implementation [39, 40]. We show a nearly significant increase in serotype 3 following PCV13 implementation in EJ (p=0.056) and no change after PCV10 implementation in PA. Whether serotype 3 will eventually decrease in a longer follow-up is yet to be seen.

The relatively stable prevalence of *S. pneumoniae* carriage despite the significant reduction in VT strains is attributed to the replacement by non-VT strains as previously described [41-45]. Serotype replacement is a universal phenomenon in which non-VT serotypes emerge and replace VT serotypes with geographic variability [42]. The emerging serotypes we observed following the two different vaccination policies were somewhat similar.

In EJ, the most notable emerging non-VT serotypes were 11A/D, 19B/C, 25/38, 40, 21, 9N/L and 12F and in PA they were 19B/C, 40, 23A and 7B/C. A study in Massachusetts, USA reported that several years following PCV7 implementation, serotypes 19A, 6A, 15B/C, 35B, and 11A emerged [46]. In Northern Japan, serotypes 15A, 23A, 11A, 10A and 35B accounted for the majority of non-VT13 serotypes after the introduction of PCV13 [47]. Serotype 6C was shown to decline following PCV13 implementation but not PCV7 or PCV10 [19, 38, 48, 49]. Interestingly, we observed emergence of 6C following PCV7 implementation, but a decrease after either PCV13 or PCV10 implementation.

Parental nasopharyngeal pneumococcal carriage was rare in our population. Carriage rates in adults were reported to be less than 10%, but higher rates were found among adults with children at home [50]. An explanation of the relatively low carriage rates we detected could be that we determined carriage via nasopharyngeal swabbing, while recent reports suggested higher yield in adults when swabbing both pharynx and nasopharynx or adding salivary testing [51-53]. While sixty percent of the carrier parents had a carrier child, only 42.7% of those parent-child co-carrier pairs had an identical serotype. Similar dis-concordance was previously reported among adults in Israel, where intra-familial transmission could not be demonstrated [54]. Children were shown to carry pneumococcal strains for months, while adults typically carry pneumococci for only very short durations [55]. Yet, this does not essentially explain the relative dis-concordance of serotypes between children and their parents. The difference between serotypes carried by children’s and their parents’ could be due to the difference of the direct PCV impact and the indirect (herd effect) impact, on their parents, particularly the potential time lag of the effects.

Our study has a few limitations, mainly due to its design as an observational study. First, vaccination policies were implemented at different times, PCV7/13 in 2009/2010 in EJ, and PCV10 in 2011 in PA. Second, while the two compared populations are closely related, they differed in several variables that were adjusted for. Third, the overall carriage in the children was relatively low, probably due to the ‘off season’ periods we chose, i.e. spring and summer, when carriage of pneumococci is lower. This was intentional in order to overcome seasonal variability, but limited the power to detect some differences. Last, this study only reports the short-term impact of the vaccines and to assess the long-term differences between the vaccines, longer follow-up is needed.

In conclusion, the unique settings of this study allowed us compare the initial effects of PCV13 and PCV10 in two closely related populations that live in two geographically proximate regions, under two different health authorities. Despite the short follow-up interval after implementation of either PCV13 in EJ or PCV10 in PA, a dramatic decrease in the VT13 serotypes (including serotypes 6A and 19A but not serotype 3) was observed. Replacement by non-VT13 was also observed in both populations regardless of the vaccination used. Longer follow-up is needed to compare the long-term effects.

## Acknowledgments

We would like to acknowledge the late Dr. Raz for taking part in establishing the PICR group. We would also like to thank Efrat Steinberger, Mulu Tiruneh, Muayyad Ghoul and Miriam Varon for coordinating the study, Aylana Reiss-Mandel, Melody Kasher, Mahmoud Ramlawi, and Julie Lowenthal for assisting in the laboratory.

The PICR members 2014: PI: Dr. Gili Regev-Yochay, Co-PI: Prof. Ziad Abdeen, Izzeldin Abullaish, Rania Abu Seir, Muhammed Affiffi, Kifaya Azmi, Yunes Bassem, Adi Cohen, Muhannad Daana, Ibrahim Dandis, Dafna Doron, Abedalla ElHamdany, Ayob Hamdan, Samantha Hasselton, Amit Hupert, Muhammed Husseini, Fuad Jaar, Laduyeh Kawather, Galia Rahav, Meir Raz, Avraham Rodity, Hector Roizin, Waeel Siag, Ora Stern, Amin Thalji, Luba Yakirevitch, Khairi Zecayra.

## Funding

This work was supported by Maccabi Healthcare Services Research Institute [MIHSR-250809] and the Israel National Institute for Health Policy Research (NIHP) [NIHP-25-10] and MERC USAid [M-33-14]. All funding sources had no involvement in the study design, collection, analysis or interpretation of the data, nor did they have a role in writing the report or in the decision to submit the paper for publication.

## Supporting information

**S1 Table**: Characteristics of the two study populations in each of the four screening periods.

